# Subgenome dominance of *ALS* target site mutations impacts herbicide resistance in allohexaploid *Echinochloa crus-galli*

**DOI:** 10.1101/2025.02.20.639371

**Authors:** Luan Cutti, Guilherme Menegol Turra, Filipi Mesquita Machado, Estéfani Sulzbach, Paula Sinigaglia Angonese, Catarine Markus, Todd Adam Gaines, Aldo Merotto

## Abstract

Herbicide target site resistance in polyploid species is more complex than diploids due to potential subgenome interactions and gene dosage of mutations. The objective of this study was to identify the level of resistance and cross-resistance patterns of *ALS* mutations located in the different subgenomes of allohexaploid *Echinochloa crus-galli*. *E. crus-galli* populations were screened and dose-response curves were performed with ALS-inhibitors from different chemical groups. The *ALS* genes of each subgenome (A, B, and C) were sequenced. Copy number variation, global relative expression, and the specific relative expression of *ALS* gene from each subgenome were performed. Out of 100 populations, 32% were resistant only to imazethapyr, and 48% were cross-resistant to all ALS-inhibitors. The mutations Ala122Thr, Ala205Asn, and Ser653Asn confer resistance only to imazethapyr, while Trp574Leu confers resistance to all four herbicides tested. *ALS* mutations were most frequent in subgenome A, but *ALS* from subgenome C had the highest expression. The biotype SAOJER-01 had Trp574Leu mutation in subgenome C and was 22 times more resistant to imazethapyr and penoxsulam than CAMAQ-01, which had the same Trp574Leu mutation in subgenome A with no differences in herbicide metabolism rate. The *ALS* genes from subgenomes A and C contribute approximately 13% and 50% respectively to the total *ALS* transcripts. The higher contribution of subgenome C to the total *ALS* transcripts pool combined with the Trp574Leu resistance mutation makes the biotype SAOJER-01 more resistant to herbicides than CAMAQ-01. This is the first study showing the effect of subgenome gene expression level on the herbicide resistance level in a polyploid weed species. In addition, while imazethapyr did not control any biotype with an *ALS* mutation, the other ALS-inhibitor herbicides were able to control biotypes carrying Ala122Thr, Ala205Asn, and Ser653Asn, which reinforces the utility of these herbicides in weed and resistance management.

## Introduction

Herbicide resistance is one of the major threats to crop protection and food safety. Weeds have been identified as resistant to several herbicide modes of action through several resistance mechanisms. The herbicide resistance mechanisms are classified into two major categories, target site and non-target site (Mucheri et al. 2024). Target site mechanisms refer to any alteration in the enzyme inhibited by the herbicide molecule that affects the binding, or the overproduction of the target enzyme due to increased expression or increased number of the coding gene across the genome (Murphy and Tranel 2019). Non-target site is a more complex character and involves different strategies utilized by plants, including increased capacity to metabolize the herbicide, usually driven by *CYP*, *UGT*, *GST*, or *AKR* genes up-regulation, and reduced translocation by vacuolar sequestration, related to ABC-transporters (Jugulam and Shyam 2019). Depending on the mechanism of resistance, cross-resistance can occur to other herbicide molecules that target the same enzyme. The Trp574Leu mutation in the *acetolactate synthase* (*ALS*) gene is well-known to confer resistance to all ALS-inhibitors in many different species (Wang et al. 2019; Gherekhloo et al. 2018); the Trp2027Cys in acetyl coenzyme-A carboxylase (ACCase) confers resistance to aryloxyphenoxypropionates and phenyl pyrazoline chemical groups, but not to cyclohexanediones from ACCase-inhibitors in *Digitaria insularis* (Takano et al. 2020). The same mutation may confer different resistance levels when compared in two different species, or even may confer different cross-resistance patterns, as shown comparing the effect of Pro197Thr in *Papaver rhoeas* and *Bromus japonicus* (Liu et al. 2024; Kaloumenos et al. 2011). The explanation for different phenotypic expression of the same allele may be related to genome biology, such as ploidy level and subgenome interactions.

Polyploidy is more frequent than diploidy in plants (te Beest et al. 2012), although this characteristic is not considered in several studies about herbicide resistance. Diploids are those with the genome composed of two sets of chromosomes of the same subgenome, while polyploids are those with three, four, five, or six chromosome sets called triploid, tetraploid, pentaploid, or hexaploid, respectively (Doyle et al. 2008; Heslop-Harrison et al. 2023). Allopolyploids are polyploids that had the genome duplication event originating from an interspecific crossing, while autopolyploids result from duplication of the same genome within a species (Doyle et al. 2008). Crops such as wheat (allotetraploid and allohexaploid varieties), oat (allohexaploid), cotton (allotetraploid), and potato (auto-allotetraploid and allopentaploid) are examples of polyploids (Akagi et al. 2022). However, the plants kingdom also includes polyploid troublesome weeds, such as *Echinochloa crus-galli* (allohexaploid), *E. colona* (allohexaploid), *E. phyllopogon* (allotetraploid) (Ye et al. 2020; Wu et al. 2022), *Poa annua* (allotetraploid) (Robbins et al. 2023), *Leptochloa chinensis* (tetraploid) (Wang et al. 2022), and *Avena fatua* (allohexaploid) (Yang et al. 1999). The consequences of polyploidization are the accumulation of redundant alleles of the same gene, making them available to new functionalization, or to combine efforts playing together on certain characteristics. When polyploidization and herbicide resistance are combined, the main consequences may be the increased number of metabolism and herbicide target coding genes, which may favor faster and more complex evolution of herbicide resistance in comparison with diploid weeds. *E. crus-galli* is an allohexaploid species, carrying two chromosome sets from three different subgenomes (AABBCC), a total of 54 chromosomes (2n=6x=54), and has been successful at invading different crop fields worldwide (Wu et al. 2022), with rapid herbicide resistance evolution to several herbicides due to target and non-target site mechanisms (Turra et al. 2023; Pan et al. 2022). The *E. crus-galli* weediness may be favored by being polyploid, for example the presence of 867 *CYP*, 227 *GST*, 361 *ABC*, and 113 *AKR* genes in its genome may favor the evolution to metabolic resistance when compared to *E. haploclada*, a diploid species with many less genes and much less agricultural impact (Ye et al. 2020; Wu et al. 2022). The herbicide target site coding genes are also amplified in polyploids; for example, in *E. crus-galli* there are three *ALS* genes (Panozzo et al. 2021) and six *ACCase* genes (Iwakami et al. 2024). The impact of polyploidization on herbicide target site resistance warrants further study.

ALS-inhibitors are herbicides utilized at very low rates that have a favorable toxicity profile and are efficient in controlling a wide range of weed species while being selective to many crops, such as rice, wheat, corn, and soybean (Zhou et al. 2007). Consequently, the ALS-inhibiting herbicide mode of action has the most resistance cases reported (Heap 2024), and the most target site resistance investigation studies. Up to date, mutations in nine *ALS* positions have been reported, including Ala122, Pro197, Ala205, Phe206, Asp376, Arg377, Trp574, Ser653, and Gly654. The most common mutations are Trp574Leu and Pro197 to a range variety of amino acids (Heap 2024; Gaines et al. 2020). The ALS-inhibiting herbicides are sorted into seven chemical groups according to HRAC (Herbicide Resistance Action Committee) 2024 classification: imidazolinones, sulfonylureas, triazolopyrimidines - type 1, triazolopyrimidines - type 2, pyrimidinylbenzoates, triazolinones, and sulfonanilides. Considering the favorable chemistry profile of ALS-inhibitors combined with the scenario where *E. crus-galli* resistant has worldwide distribution in crops with limited options for crop selective herbicides, the knowledge of cross-resistance provided by target site mutations could lead to a rationale rotation with different chemical groups from ALS-inhibitors, helping to utilize this valuable mode of action. In addition, diving into the weed genomics era and the knowledge about the presence of different alleles in polyploid species must be considered when studying target site mutations. The current effort to generate genomic resources for weed species (Montgomery et al. 2024) and a deeper understanding of genome biology raises the question about the unknown effects of subgenome dominance on expression of target site coding genes in polyploid weeds and consequences for herbicide resistance. The objective of this study was to identify the level of resistance and cross-resistance patterns of *ALS* mutations located in the different subgenomes of *E. crus-galli*.

## Material and Methods

### Population collections and ALS-inhibitor screening

Seeds from 100 *E. crus-galli* populations were collected in Southern Brazil, mainly Rio Grande do Sul state between 2011-2020. The seeds were germinated and at 3-4 leaf stage screened with four active ingredients from different chemical groups of ALS-inhibitors herbicides: imazethapyr (imidazolinones), penoxsulam (triazolopyrimidine – type 2), bispyribac-sodium (pyrimidinylbenzoates), and nicosulfuron (sulfonylureas). The doses sprayed were the label rate of each herbicide, 106 g ha^-1^ of imazethapyr (Imazetapir Plus Nortox, Nortox, 106 g L^-1^) + adjuvant Dash (BASF) 0.5% v/v, 60 g ha^-1^ of penoxsulam (Ricer, Corteva, 240 g L^-1^) + adjuvant Veget’Oil (Oxiquímica, 0.5% v/v), 50 g ha^-1^ of bispyribac-sodium (Nominee 400 SC, Iharabras, 400 g L^-1^) + adjuvant Dash (BASF) 0.5% v/v, and 60 g ha^-1^ of nicosulfuron (Nicosulfuron Nortox 40 SC, Nortox, 40 g L^-1^), at spray volume of 200 L ha^-1^. The application was performed with an automated spray chamber (Generation III, Devries manufacturing). The screening was conducted in a greenhouse (28°C ± 5, supplemented with 500 to 600 μmol m^−2^ s^−1^ in a 14/10h (light/dark) cycle) and the plants were kept flooded during the duration of the experiment. Each treatment had four replicates. The control efficacy was assessed at 28 days after treatment (DAT), where 0% indicated no injury and 100% indicated dead plants. Plants with up to 80% injury while remaining alive were considered resistant.

### Dose-response curves to four ALS-inhibitor chemical groups

Eight random *E. crus-galli* populations showing different cross-resistance patterns from the previous experiment, and two susceptible populations were selected to perform dose-response curves. These selected populations had *ALS* gene mutations that were sequenced in a previous study (Turra et al. 2023), with one population carrying Ala122Thr (BAGE-01), one with Ala205Asn (SANTPAT-01), three with Trp574Leu (CAMAQ-01, CAPV-03, and SAOJER-01), and three with Ser653Asn (ARRGR-01, PALMS-01, and 423). These populations had been self-pollinated for four generations to ensure homozygosity, and these inbred lines are referred to as biotypes. The herbicides sprayed were imazethapyr, penoxsulam, bispyribac-sodium, and nicosulfuron, utilizing the rates described previously as reference label rates. The doses sprayed are described in Suppl. Table 1. The experiment parameters and spray conditions were the same as described above. Dry biomass was assessed at 28 DAT. The data were submitted to analysis of variation (ANOVA) and when the interaction between factors was significant (p ≤ 0.05), the averages were fitted to the three-parameter log-logistic non-linear regression model using the *drc* package in R (Ritz and Streibig, 2005) as follows: y = d/(1+(x/e)^b); where y is the dry biomass, x is the herbicide dose, b is the curve slope at the inflection point, d is the upper limit, and e is the inflection point, representing the dose that reduced 50% of the dry biomass (GR50 parameter). The latest was utilized to compare the resistance level of each biotype.

### Dose-response curve with metabolic inhibitors

Dose-response curves to penoxsulam in combination with the P450 inhibitor malathion (Oliveira et al. 2018) or the glutathione-S-transferase (GST) inhibitor NBD-Cl (4-Chloro-7-nitrobenzofurazan) (Cummins et al. 2013) were performed utilizing two *E. crus-galli* biotypes carrying Trp574Leu mutation CAMAQ-01 and SAOJER-01, and one susceptible MOSTS-01. The penoxsulam doses were 0; 20; 60; 180; 540; 1,620; 4,860; and 14,580 g ha^-1^ for CAMAQ-01, while 0; 60; 180; 540; 1,620; 4,860; 14,580; and 43,740 g ha^-1^ for the SAOJER-01, and 0; 0.93; 1.87; 3.75; 7.5; 15; 30; 60 g ha^-1^ for the susceptible MOSTS-01, plus adjuvant Veget’Oil 0.5% v/v. Malathion was sprayed 2 hours before the herbicide at 1,300 g ha^-1^, while NBD-CL was sprayed 48 hours before at 270 g ha^-1^ diluted in a 1:1 water:acetone solution. The experiment procedures, conditions, and data analysis followed the same described previously.

### Copy number variation

The DNA of the eight resistant biotypes and one susceptible was extracted following the CTAB method with some adaptations (Doyle and Doyle 1987). The *ALS* copy number variation was assessed utilizing a primer pair that targets the three *ALS* genes of *E. crus-galli* (Table 1), and the housekeeping gene utilized was *GAPDH* (*glyceraldehyde-3-phosphate dehydrogenase*) (Suppl. Table 2). The master mix used was Sso Advanced Universal SYBR® Green Supermix (Bio-Rad) following the manufacturer protocol instructions, 98°C for 3 min and 40 cycles of 98°C for 15s and ‘annealing temperature’ for 30s, followed by one step of 65°C for 5s and 95°C for 50s, utilizing the equipment CFX96 Real-time System (Bio-Rad). The copy number variation was calculated relative to the susceptible biotype following the method 2^-ΔΔCt^ (Dussault and Pouliot 2006). Each biotype had four biological replicates and three technical replicates. The data were compared according to the ANOVA and when significant (p ≤ 0.05) the averages were compared according to Tukey 5%.

**Table 1.**
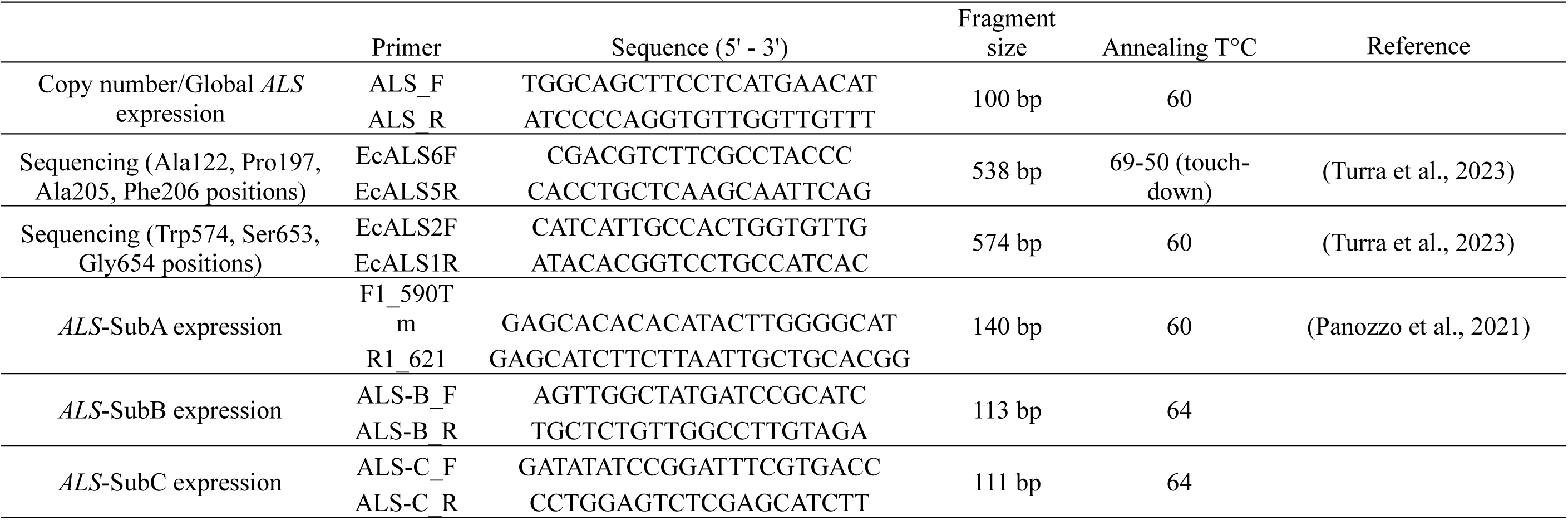
*ALS* primers utilized for partial gene sequencing, copy number variation, global *ALS* relative expression, and specific *ALS* relative expression of subgenomes A, B, or C.

### Subgenome location of *ALS* mutations

To identify possible target site mutations, the DNA of three individuals from 19 populations from the initial screening experiment had partial *ALS* gene amplified utilizing the primers targeting the positions with the mutations previously identified (Table 1) following the conditions described in Turra *et al*., (2023). These primers target the *ALS* gene of all three *E. crus-galli* subgenomes. To sequence the individual *ALS* genes of each subgenome, the PCR product of one individual of each biotype was inserted into a vector and then transformed in *E. coli* utilizing TOPO™ TA Cloning™ Kit for Sequencing, with One Shot™ TOP10 Chemically Competent *E. coli* (Invitrogen), following the manufacturer’s protocol. The plates with colonies were incubated for 24h at 37°C. Twelve individual colonies of a single individual of each biotype were isolated to perform a colony-PCR utilizing the primers targeting the position where the mutation was previously identified, and the PCR product was Sanger sequenced. The distinction of the three *ALS* genes was based on single nucleotide polymorphisms (SNPs) between them, comparing the Genbank sequences MH013494.1 (*ALS*-SubA), MG188321.1 (*ALS*-SubB), and MH013493.1 (*ALS*-SubC) (Suppl. Fig. 1) and the twelve colonies sequenced of each individual from each biotype. The subgenome was attributed to each sequence comparing the three Genbank sequences with the *ALS* genes of the last version of *E. crus-galli* genome (Wu et al., 2022). The absence of double peaks in the SNPs position that distinguish the subgenomes indicated a single *ALS* amplification.

### Expression of *ALS* of each subgenome

Seven *E. crus-galli* biotypes, one susceptible (MOSTS-01), three with the *ALS* mutation Trp574Leu (CAMAQ-01, CAPV-03, and SAOJER-01), and three with Ser653Asn (ARRGR-01, PALMS-01, and 423) had leaf tissue collected from untreated plants and 48 hours after penoxsulam treatment at 72 g ha^-1^, plus adjuvant MSO (methylated seed oil) 0.5% v/v, and spray volume of 200 L ha^-1^. The plants were grown in a growth chamber (14h light/10h dark, temperature 30 °C day/25 °C night), and were sprayed with an automated spray chamber. The tissue of the two youngest leaves was immediately frozen in liquid nitrogen after collection, and the RNA was extracted utilizing the Direct-zol RNA Miniprep Plus kit (ZymoResearch). The cDNA synthesis was performed using the iScript cDNA Synthesis kit (Bio-Rad). The expression of the *ALS* from each subgenome was assessed utilizing specific primers targeting SNPs that differentiate them (Fig. 1A). The primer pairs targeting specifically the *ALS* from subgenomes B and C were designed utilizing the Primer3Plus software (https://www.bioinformatics.nl/cgi-bin/primer3plus/primer3plus.cgi) (Table 1), while the primer pair targeting *ALS* from subgenome A was obtained from Panozzo *et al*. (2021) (Table 1). The specificity of our primers is shown by the absence of double peaks at the position that differentiates the subgenomes (Fig. 1B). The primer pair targeting all three *ALS* genes was the same utilized for copy number variation. The housekeeping genes utilized were *eIF4B1* (e*ukaryotic translation initiation factor 4B1*), *GAPDH*, *18S* (*18S ribosomal RNA*), *28S* (2*8S ribosomal RNA*), and *RUB* (*rubisco*) (Suppl. Table 2), which are common control genes utilized in herbicide qPCR studies (Duhoux and Délye 2013; Chen et al. 2017; Wrzesińska et al. 2016; Liu et al. 2019). The *eIF4B1* gene was identified as the most stable according to the stability analysis using the algorithm Normfinder in the RefFinder platform. The qPCR was performed following the protocol described for the ‘copy number variation’, with four biological and three technical replicates. The relative expression was calculated according to the method 2^-ΔΔCt^ (Dussault and Pouliot 2006), utilizing the *ALS* from subgenome A of each biotype as the reference. The data were compared according to ANOVA and when significant (p ≤ 0.05) the averages were compared according to Tukey 5%. Based on the relative expression results, we estimated the percentage contribution of each *ALS* subgenome to the total amount of *ALS* transcripts.

**Fig. 1.**
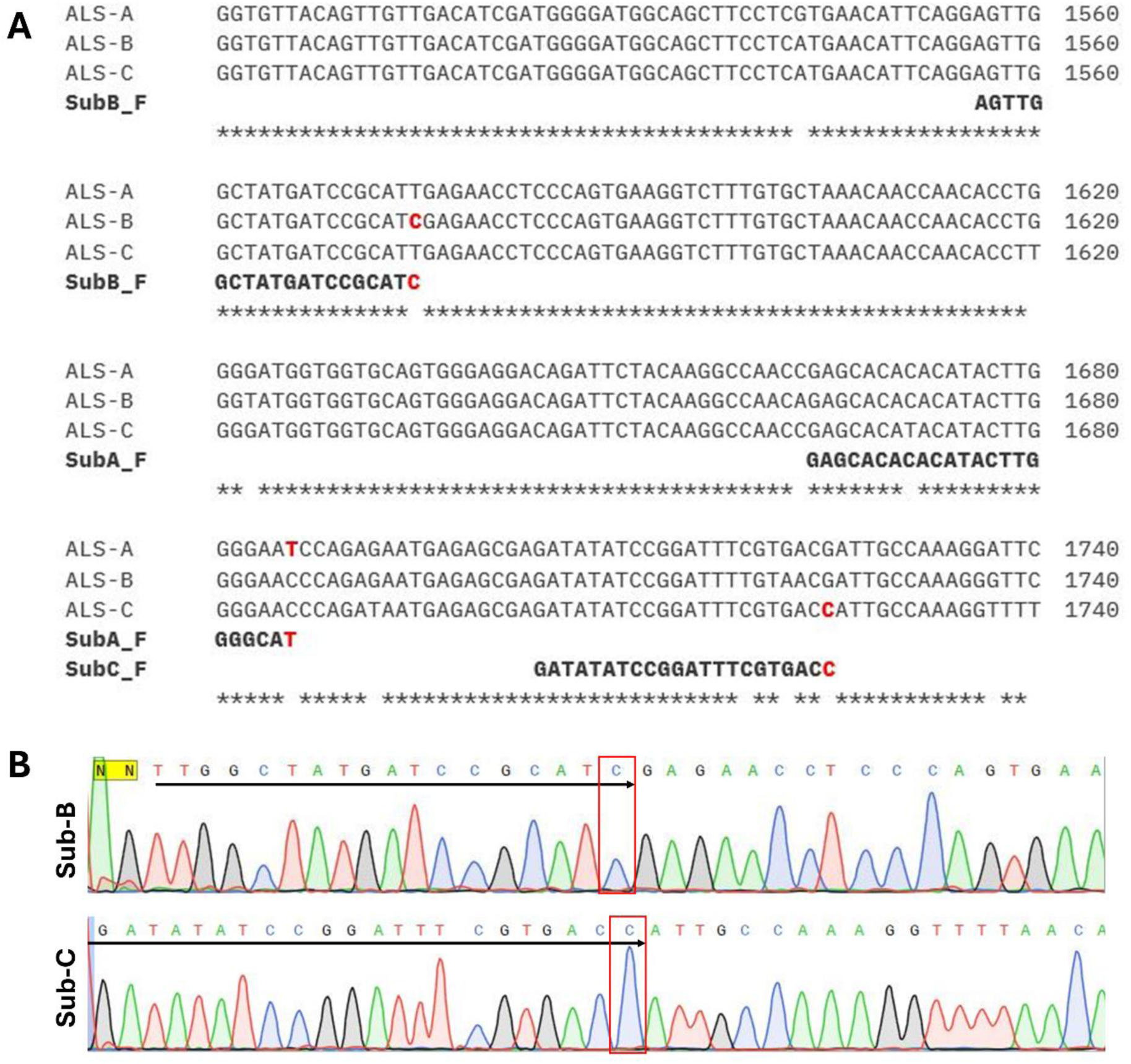
Confirmation of specificity of the primers targeting the three *ALS* from different subgenomes. (A) Partial *E. crus-galli ALS* sequence showing the SNPs that differentiate them in each subgenome, and the forward primers targeting the SNPs at the 3’-end. (B) Sequencing of the PCR product when utilizing the *ALS*-specific primers targeting subgenome B or C, showing the absence of double peak at the primer 3’-end where the specific SNPs are.

## Results

*E. crus-galli* populations resistant to ALS inhibitors are spread across Southern Brazil, mainly in Rio Grande do Sul state (Fig. 2A). Most of the populations evaluated, 48%, showed cross-resistance to imazethapyr, penoxsulam, bispyribac-sodium, and nicosulfuron, while 32% were resistant only to imazethapyr and 20% were still susceptible to all ALS-inhibitors tested (Fig. 2B; Suppl. Table 3). The western region of the state had more prevalence of cross-resistant populations, while the eastern region had more imazethapyr single-resistant and susceptible populations. All populations resistant to ALS inhibitors were resistant to imazethapyr (80%) (Fig. 2B).

**Fig. 2.**
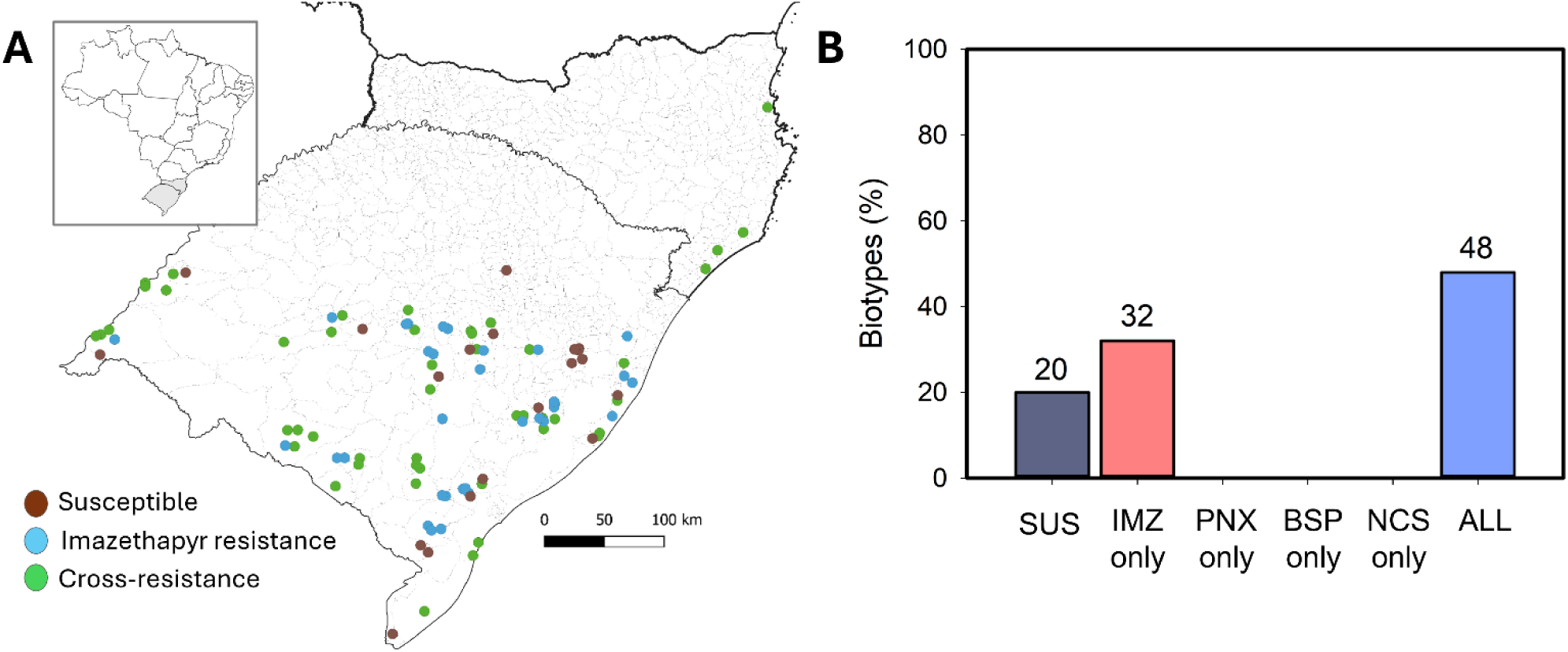
Mapping the resistance status of 100 *Echinochloa* biotypes in Southern Brazil, and cross-resistance characterization. (A) Distribution of *Echinochloa* biotypes susceptible and resistant to ALS-inhibitor herbicides in Southern Brazil. (B) Percentage of biotypes characterized as susceptible, resistant to only one ALS-inhibitor, or with cross-resistance to all four ALS-inhibitors tested.

Four *ALS* mutations Ala122Thr, Ala205Asn, Trp574Leu, and Ser653Asn were identified in *E. crus-galli* in a previous study (Turra et al. 2023). All the mutations conferred resistance to imazethapyr but at different levels. The mutations Ala122Thr and Ala205Asn conferred intermediate imazethapyr resistance levels when compared to Trp574Leu (usually the highest) and Ser653Ans (usually the lowest) (Table 2). However, variation between the biotypes carrying the same mutation was observed, and no clear resistance level could be linked to each mutation. The biotype CAMAQ-01 with Trp574Leu had the smallest imazethapyr resistance factor (RF), while SAOJER-01 also with Trp574Leu had the highest RF among all populations evaluated. The SAOJER-01 was 3.3 times more resistant to imazethapyr than CAPV-03, and 22.3 times more than CAMAQ-01, all three with Trp574Leu. A clear RF difference between biotypes carrying the same mutation Ser653Asn was also observed. The biotype 423 is 6.4 and 7.8 times more resistant to imazethapyr than ARRGR-01 and PALMS-01, all three with Ser653Asn (Table 2). Penoxsulam, bispyribac-sodium, and nicosulfuron resistance were observed only for biotypes carrying the Trp574Leu mutation (Table 2). Notably, biotypes with mutations Ala122Thr, Ala205Asn, and Ser653Asn also had higher RF to bispyribac-sodium, nicosulfuron, and mainly penoxsulam when compared to the susceptible biotype; however, they were not considered resistant because they did not survive the herbicide label rates. The differential resistance levels between the three populations carrying Trp574Leu populations were observed for all four ALS-inhibitors tested (Table 2), as shown for penoxsulam in Fig. 3.

**Fig. 3.**
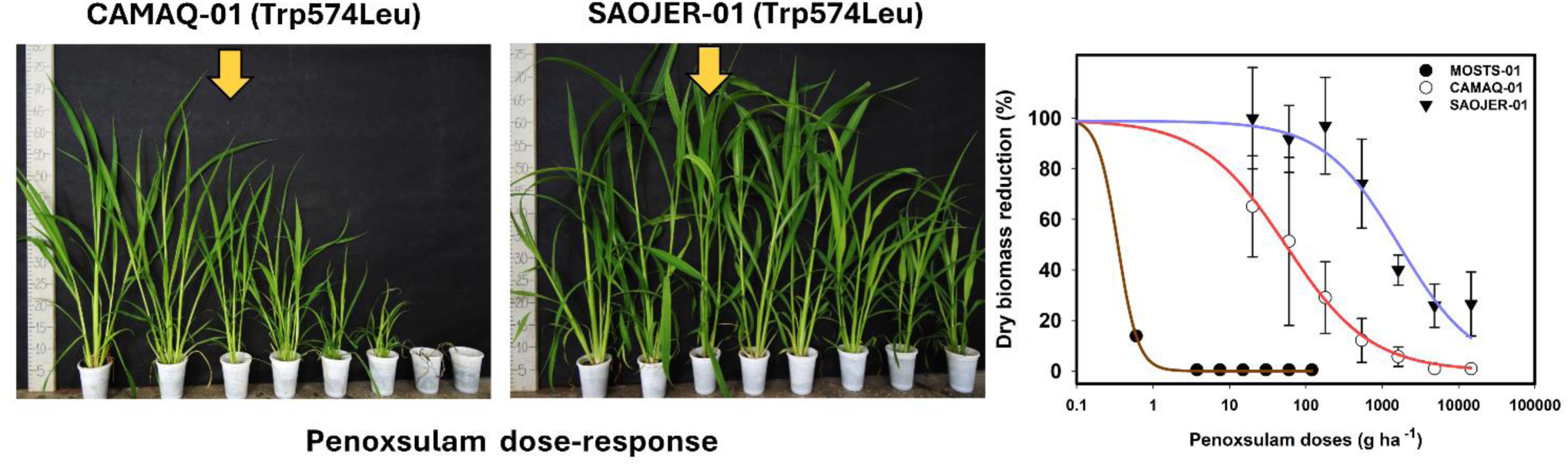
Differential resistance level to penoxsulam of two *E. crus-galli* biotypes carrying the same Trp574Leu mutation in *ALS* gene. Yellow arrow indicates the label rate (60 g ha^-1^).

**Table 2.** Dose-response curve parameters to four ALS inhibitors (imazethapyr, penoxsualm, bispyribac-sodium, nicosulfuron) from different chemical groups (imidazolinones, triazolopyrimidine – type 2, pyrimidinylbenzoates, and sulfonylureas) utilizing *E. crus-galli* biotypes with four different *ALS* mutations.

| Biotypes | Mutation | b | d | e(GR50) | GR50 Lower | GR50 Upper | RF |
| --- | --- | --- | --- | --- | --- | --- | --- |
| Imazethapyr |  |  |  |  |  |  |  |
| MOSTS-01 | No | 3.53* | 4.23* | 2.85* | 2.19 | 3.51 | - |
| CAPL-01 | No | 3.69* | 3.48* | 2.80* | 1.90 | 3.70 | 1.0 |
| BAGE-01 | Ala122Thr | 1.27* | 4.78* | 82.65* | 60.20 | 105.11 | 29.0 |
| SANTPAT-01 | Ala205Asn | 1.55* | 5.11* | 92.60* | 72.82 | 112.39 | 32.5 |
| CAMAQ-01 | Trp574Leu | 0.53* | 3.06* | 20.18<sup>ns</sup> | -11.52 | 51.89 | 7.1 |
| CAPV-03 | Trp574Leu | 0.50* | 3.79* | 134.31* | 32.87 | 235.75 | 47.1 |
| SAOJER-01 | Trp574Leu | 0.58* | 2.90* | 449.87* | 82.33 | 817.41 | 157.8 |
| ARRGR-01 | Ser653Asn | 1.54* | 4.83* | 43.72* | 34.22 | 53.23 | 15.3 |
| PALMS-01 | Ser653Asn | 0.97* | 6.27* | 35.92* | 27.22 | 44.62 | 12.6 |
| 423 | Ser653Asn | 2.14* | 2.65* | 279.05* | 183.56 | 374.55 | 97.9 |
| Penoxsulam |  |  |  |  |  |  |  |
| MOSTS-01 | No | 0.93<sup>ns</sup> | 5.98* | 0.04<sup>ns</sup> | -0.14 | 0.23 | - |
| CAPL-01 | No | 1.11* | 6.65* | 0.15<sup>ns</sup> | -0.04 | 0.35 | 3.8 |
| BAGE-01 | Ala122Thr | 3.15* | 3.15* | 4.01* | 3.02 | 5.01 | 100.3 |
| SANTPAT-01 | Ala205Asn | 0.75* | 4.68* | 9.50* | 4.77 | 14.24 | 237.6 |
| CAMAQ-01 | Trp574Leu | 0.78* | 3.27* | 54.78* | 12.50 | 97.05 | 1369.4 |
| CAPV-03 | Trp574Leu | 0.68* | 3.12* | 168.76* | 34.27 | 303.25 | 4219.1 |
| SAOJER-01 | Trp574Leu | 0.79* | 4.14* | 1231.32* | 597.30 | 1865.34 | 30783.0 |
| ARRGR-01 | Ser653Asn | 4.13<sup>ns</sup> | 4.72* | 2.40<sup>ns</sup> | -1.02 | 5.81 | 60.0 |
| PALMS-01 | Ser653Asn | 4.99<sup>ns</sup> | 3.44* | 2.82<sup>ns</sup> | -2.32 | 7.95 | 70.5 |
| 423 | Ser653Asn | 2.61<sup>ns</sup> | 2.99* | 2.29<sup>ns</sup> | -1.40 | 5.98 | 57.3 |
| Bispyribac-sodium |  |  |  |  |  |  |  |
| MOSTS-01 | No | 1.89* | 6.78* | 0.63* | 0.45 | 0.80 | - |
| CAPL-01 | No | 1.96* | 3.92* | 1.27* | 0.87 | 1.67 | 2.0 |
| BAGE-01 | Ala122Thr | 3.97* | 5.42* | 3.51* | 1.42 | 5.60 | 5.6 |
| SANTPAT-01 | Ala205Asn | 2.60* | 5.89* | 7.61* | 6.55 | 8.67 | 12.1 |
| CAMAQ-01 | Trp574Leu | 0.77* | 6.03* | 20.60* | 12.79 | 28.41 | 32.8 |
| CAPV-03 | Trp574Leu | 0.92* | 2.91* | 83.13* | 43.96 | 122.31 | 132.2 |
| SAOJER-01 | Trp574Leu | 0.89* | 5.43* | 167.34* | 98.95 | 235.72 | 266.1 |
| ARRGR-01 | Ser653Asn | 3.15* | 4.62* | 4.77* | 3.84 | 5.70 | 7.6 |
| PALMS-01 | Ser653Asn | 3.44<sup>ns</sup> | 6.15* | 3.20<sup>ns</sup> | -4.08 | 10.48 | 5.1 |
| 423 | Ser653Asn | 1.53* | 3.25* | 4.99* | 3.28 | 6.69 | 7.9 |
| Nicosulfuron |  |  |  |  |  |  |  |
| MOSTS-01 | No | 1.88* | 6.01* | 4.48* | 3.42 | 5.54 | - |
| CAPL-01 | No | 3.67* | 4.02* | 12.11* | 10.05 | 14.17 | 2.7 |
| BAGE-01 | Ala122Thr | 5.07* | 4.60* | 8.56* | 6.71 | 10.42 | 1.9 |
| SANTPAT-01 | Ala205Asn | 2.54* | 7.69* | 17.38* | 14.16 | 20.61 | 3.9 |
| CAMAQ-01 | Trp574Leu | 1.34* | 6.79* | 36.05* | 27.04 | 45.07 | 8.1 |
| CAPV-03 | Trp574Leu | 1.55* | 3.37* | 81.23* | 55.15 | 107.31 | 18.1 |
| SAOJER-01 | Trp574Leu | 1.28* | 5.48* | 155.99* | 104.26 | 207.73 | 34.8 |
| ARRGR-01 | Ser653Asn | 5.78<sup>ns</sup> | 4.58* | 14.74* | 13.31 | 16.17 | 3.3 |
| PALMS-01 | Ser653Asn | 10.16<sup>ns</sup> | 4.77* | 15.91* | 9.17 | 22.66 | 3.6 |
| 423 | Ser653Asn | 3.36* | 2.56* | 24.51* | 19.62 | 29.39 | 5.5 |

\*significant parameter (p<0.05); <sup>ns</sup>non-significant parameter.

We investigated the location of *ALS* mutations in the allohexaploid *E. crus-galli* genome. The *ALS* gene is located in Chr07 of subgenomes A, B, and C. Sixteen populations carrying Ala122Thr, Ala205Asn, Trp574Leu, or Ser653Asn had these mutations in *ALS* from subgenome A, while one biotype had a Trp574Leu in subgenome C, and one had a double-mutation with Trp574Leu in subgenome A + Ala122Asn in subgenome C (Table 3). The biotype with the highest RF, SAOJER-01, had the Trp574Leu mutation in subgenome C, while CAPV-03 and CAMAQ-01, with smaller RF, had the same mutation in subgenome A. All three biotypes with Ser653Asn had the mutation in subgenome A. We moved forward with additional experiments to understand the RF differences between the contrasting biotypes SAOJER-01 and CAMAQ-01 carrying Trp574Leu on different subgenomes focusing on the investigation of metabolic resistance. No effect on resistance occurred with the GST inhibitor, but a strong P450 inhibitor effect was observed in both CAMAQ-01 and SAOJER-01 biotypes sprayed with penoxsulam, with an approximately 12-fold reduction in GR_50_ in CAMAQ-01 and an approximately 7-fold reduction in GR_50_ in SAOJER-01 due to the P450 inhibitor application (Table 4). This result indicates that both biotypes have the P450 metabolic mechanism acting on herbicide detoxification in addition to the target site resistance mutation. In addition, the RF to penoxsulam (without inhibitor) for both CAMAQ-01 and SAOJER-01 in this second experiment (Table 4) was higher than the previous experiment (Table 2), indicative of an environmental effect on phenotypic resistance level that is common with metabolic resistance (Gaines et al. 2020). Note that the penoxsulam doses sprayed are different in both experiments, resulting in slight variations in the estimation of dose-response parameters. However, SAOJER-01 was still much more resistant than CAMAQ-01 when treated with the P450 inhibitor, which suggests some additional mechanism to explain the difference in resistance level between them.

**Table 3.**
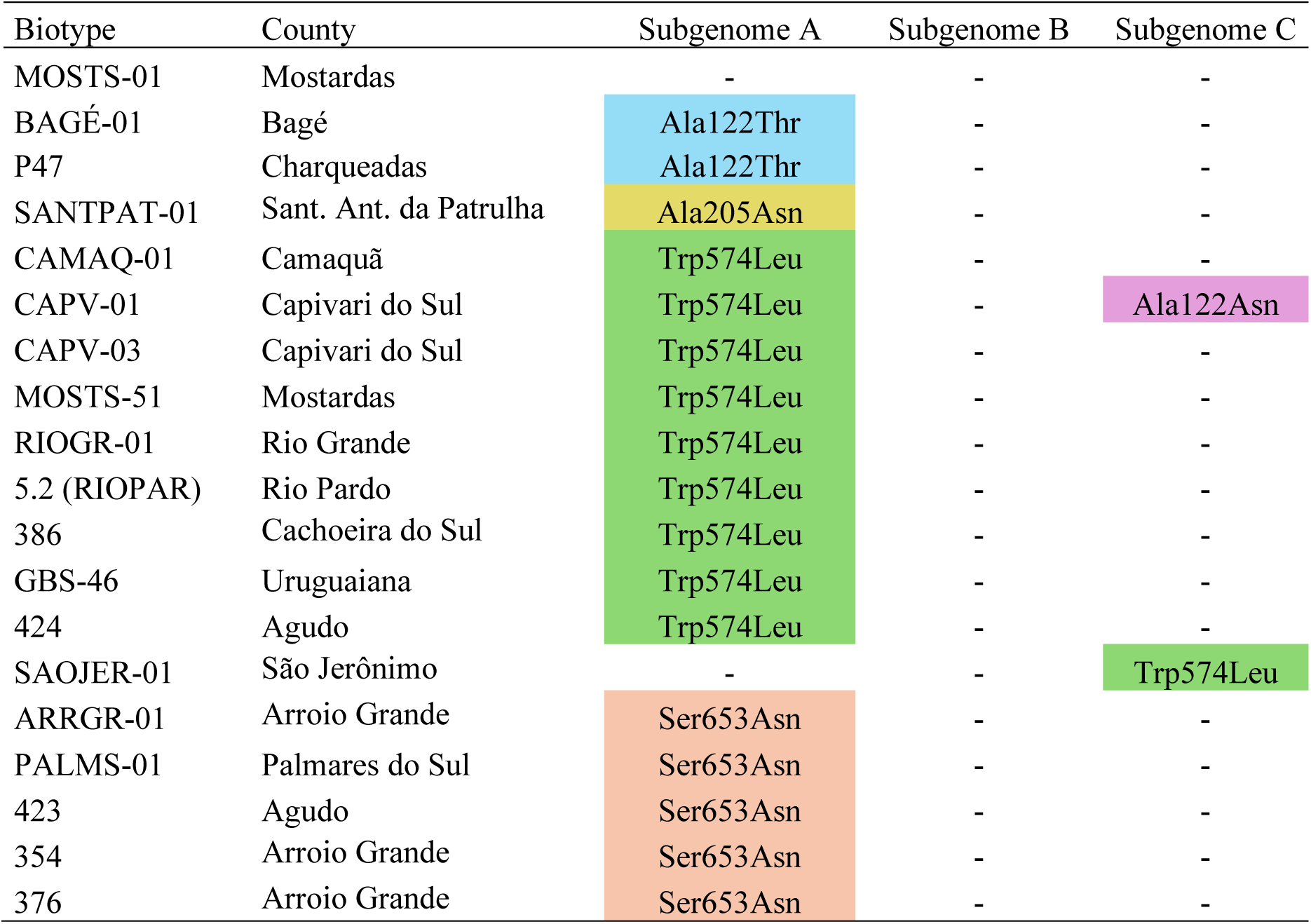
*ALS* mutations and their subgenome location in 18 *E. crus-galli* biotypes resistant to ALS-inhibitors from different counties in South Brazil.

**Table 4.** Penoxsualm dose-response curve investigating the presence of metabolic resistance in two resistant *E. crus-galli* biotypes with the same Trp574Leu mutation but showing different resistance levels, and one susceptible. Malathion and NBD-Cl were the P450 and GST enzyme metabolic inhibitors, respectively.

| Biotypes | Inhibitors | *b* | *d* | *e* (*GR*<sub>50</sub>) | *GR*<sub>50</sub> Lower | *GR*<sub>50</sub> Upper | RF<sup>1</sup> |
| --- | --- | --- | --- | --- | --- | --- | --- |
| MOSTS-01 | No | 5.34<sup>ns</sup> | 3.87* | 0.77<sup>ns</sup> | 0.42 | 1.11 | 1.00 |
|  | P450 | 3.80<sup>ns</sup> | 4.17* | 0.55<sup>ns</sup> | -0.22 | 1.32 | 0.71 |
|  | GST | 4.71<sup>ns</sup> | 3.60* | 0.68* | 0.04 | 1.32 | 0.88 |
| CAMAQ-01 | No | 0.47* | 3.81* | 132.94* | 59.94 | 205.93 | 1.00 |
|  | P450 | 0.33* | 3.16* | 11.16<sup>ns</sup> | -4.01 | 26.33 | 0.08 |
|  | GST | 0.34* | 3.04* | 140.07* | 7.21 | 272.92 | 1.05 |
| SAOJER-01 | No | 0.35* | 3.05* | >43,740<sup>ns</sup> | - | - | 1.00 |
|  | P450 | 0.26* | 3.57* | 6,032.30* | 447.11 | 11617.55 | <0.14 |
|  | GST | 0.34* | 2.98* | >43,740<sup>ns</sup> | - | - | 1.00 |

<sup>1</sup>Resistance Factor (RF) compares the *GR*<sub>50</sub> of metabolic inhibitors with that of no metabolic inhibitor in each biotype;
\*significant parameter (p<0.05); <sup>ns</sup>non-significant parameter.

No differences in copy number (Fig. 4A) or differences in global *ALS* expression (targeting all three subgenomes) (Fig. 4B) were observed to explain the different RF levels between the susceptible and resistant biotypes. Two biotypes (BAGE-01 and 423) showed half the number of *ALS* copies compared to the susceptible control, but they were not further investigated in this study. However, differential *ALS* expression from each subgenome in each population was observed. All populations carrying Trp574Leu, Ser653Asn, and susceptible had higher expression of *ALS* from subgenome C and B than A, except ARRGR-01 in the absence of penoxsulam (Fig. 4C). After spraying penoxsulam, the expression of *ALS* from subgenome C is the one with higher expression while the *ALS* from subgenome A was the least expressed in penoxsulam resistant populations with Trp574Leu mutation (Fig. 4D). The populations with Ser653Asn and susceptible showed a different pattern of *ALS* subgenome expression after penoxsulam treatment (Fig. 4D). These biotypes were susceptible to penoxsulam, and the expression of *ALS* from each subgenome at 48 hours after herbicide spray may be a consequence of cellular death. Utilizing the relative expression results, we estimated the contribution of *ALS* from each subgenome to the total amount of *ALS* transcripts (Fig. 4E; Fig. 4F). In untreated samples, the *ALS* from subgenome C contributed approximately 46-47% of the total *ALS* transcripts, while the *ALS* from subgenome A contributed about 13-15% in populations with Trp574Leu mutation (Fig. 4E). After penoxsulam spray, the *ALS* from subgenome C contributed 47-50%, while subgenome A contributed 12-15% (Fig. 4F). SAOJER-1 is the most herbicide-resistant biotype and also the only biotype with a Trp574Leu mutation in subgenome C, which is the most expressed subgenome. The least resistant biotypes are those with the Trp574Leu mutation in subgenome A, which is the least expressed subgenome (Table 2; Table 3; Fig. 4C; Fig. 4D). The higher quantity of mutant transcripts in the total pool of *ALS* transcripts in SAOJER-01 makes it more resistant. The highest penoxsulam dose tested in our experiment, 43,740 g ha^-1^ (which represents 729x the label rate), was not able to kill the SAOJER-01 plants (Table 4).

**Fig. 4.**
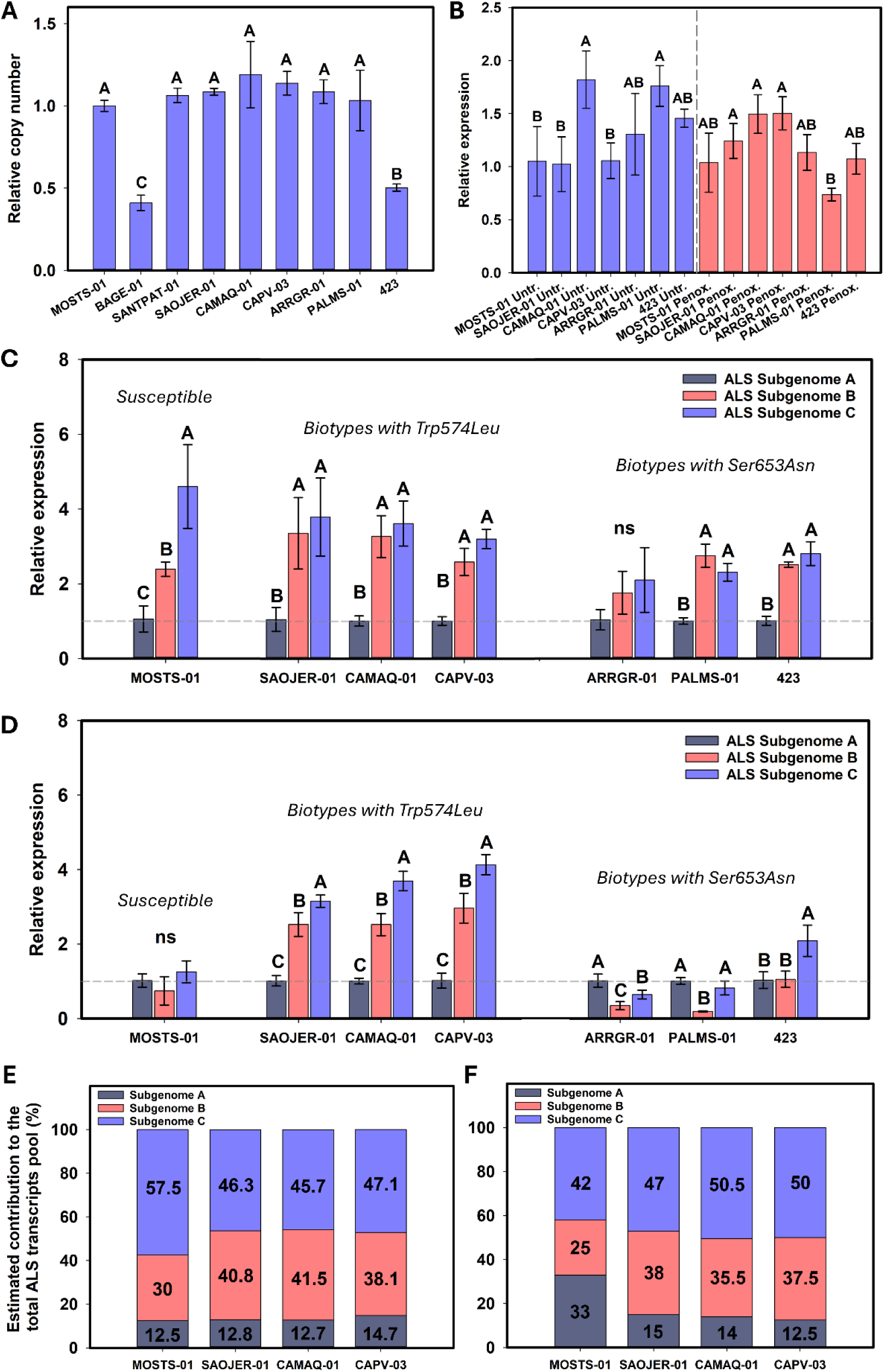
(A) *ALS* copy number variation analysis in *E. crus-galli* biotypes showing different *ALS* mutations. (B) Relative expression of total *ALS* (targeting the three subgenomes) in untreated and treated with penoxsulam. (C) Relative expression of *ALS*-SubA, *ALS*-SubB, and *ALS*-SubC in one susceptible biotype, three with Trp574Leu mutation, and three with Ser653Asn mutation without herbicide application. (D) Relative expression of *ALS*-SubA, *ALS*-SubB, and *ALS*-SubC in the same biotypes previously described 48h after penoxsulam application. (E) Estimated contribution based on relative expression analysis of each subgenome to the total *ALS* transcripts pool in one susceptible and three biotypes with Trp574Leu mutation untreated. (F) Estimated contribution based on relative expression analysis of each subgenome to the total *ALS* transcripts pool in one susceptible and three biotypes with Trp574Leu mutation 48h after penoxsulam application.

## Discussion

Besides reporting the *ALS* mutations in *E. crus-galli* and the different cross-resistance patterns, we identified that they are prevalent in subgenome A in *E. crus-galli*. Only the biotypes SAOJER-01 and CAPV-01 had a mutation in *ALS* from subgenome C, Trp574Leu and Ala122Asn, respectively. The occurrence of mutations in all *E. crus-galli ALS* subgenomes has been reported (Löbmann et al. 2021; Yang et al. 2021). However, our study is the first one sequencing multiple hexaploid biotypes to evaluate in which subgenome *ALS* mutations are prevalent. The biotype SAOJER-01 with the mutation in subgenome C was 22 times more resistant to imazethapyr and penoxsulam than CAMAQ-01 with the same mutation but in subgenome A. Biotypes with the Ser653Asn mutation in subgenome A also showed different resistance levels but were not further explored in this study. P450-mediated herbicide metabolism was present in both SAOJER-01 and CAMAQ-01 and the resistance difference between the two biotypes carrying the same Trp574Leu mutation is probably not only explained by metabolism. However, a differential expression level of the *ALS* genes from the three subgenomes was identified, with the *ALS* gene in subgenomes B and C the most expressed, and the one in subgenome A least expressed in all biotypes analyzed, except ARRGR-01. The up-regulation of *ALS* from subgenome C with the Trp574Leu mutation increased dosage of the resistant allele, causing most of the transcripts in the pool to be resistant. Another study shows *E. oryzicola* with the *ALS* mutation Trp574Leu in *ALS1* (the *ALS* from subgenome A) (Panozzo et al. 2021), which corroborates our results, where most mutations were also in subgenome A. However, their findings show higher expression of *ALS1* from subgenome A than subgenome B (*E. oryzicola* is tetraploid), while in our results the subgenome A, in hexaploid *E. crus-galli*, was the least expressed in the resistant biotypes.

Corroborating our results, the analysis of the differential expression of *E. crus-galli* subgenomes revealed that a significantly higher proportion of suppressed genes occurred in subgenome A than in subgenomes B and C (Ye et al. 2020). The preference to express a gene or multiple genes from one subgenome than another is called subgenome dominance, which is an underlying mechanism in allopolyploid species, but not in autopolyploids (Bird et al. 2018; Alger and Edger 2020). A single subgenome may play a major role in the phenotype of allopolyploids (Alger and Edger 2020). The dominance of a gene from one particular subgenome regarding the herbicide resistance phenotype does not mean that all genes from this subgenome will dominate all the other characteristics. For example, in tetraploid *T. turgidum*, some developmental traits are controlled by the A subgenome, while responses to stress and diseases are controlled by the B subgenome (Feldman et al. 2012).

Differential herbicide resistance according to the subgenome location of the mutation has been reported in a few species, but rarely or never evaluating the subgenome differential expression and herbicide resistance. In wheat (*Triticum aestivum*), the *ALS* Ser653Asn mutation in subgenome D conferred higher resistance to imazethapyr than in subgenome A or B (Chen et al. 2021). Also in wheat, the *ACCase* Ala2004Val mutation in subgenome A conferred higher resistance to quizalofop than in subgenome D or B (Ostlie et al. 2015). In *Sonchus oleraceus*, a tetraploid species, the presence of Pro197Ser in *ALS* from different subgenomes did not confer differential resistance to metsulfuron-methyl or imazamox + imazethapyr (Merriam et al. 2023). In *Monochoria vaginalis*, a tetraploid species, the presence of Pro197Ser or Asp376Glu in *ALS1* or *ALS3* did not confer differential resistance to imazosulfuron. In this case, the expression of *ALS1* and *ALS3* are similar and higher than the other three *ALS* genes found, but it is unknown if they are from the same subgenome or not (Tanigaki et al. 2021). In general, the location of mutations in one subgenome is sufficient to result in herbicide-resistant phenotype, and it does not matter if the subgenome has higher or lower expression than the others. However, in the case of mutations that confer low-resistance level, for example, Ser653Asn to imazethapyr in *E. crus-galli*, the location in a lower expressed subgenome has a practical management consequence, as increasing the herbicide dose may be sufficient to control it. However, in the case of Trp574Leu mutation, this concept does not apply. Conversely, the presence of mutations in a lower expressed subgenome in a crop species, such as wheat, can result in a higher risk of injuries and decreased yield. It is important to highlight that the location of the same mutation in different subgenomes does not impact the cross-resistance pattern.

The screening of 100 populations collected in Southern Brazil shows the distribution of ALS-inhibitors resistance in *E. crus-galli*. All resistant populations were resistant to imazethapyr; however, not all imazethapyr-resistant were resistant to penoxsulam, bispyribac-sodium, and nicosulfuron. In addition, when the population showed resistance to one of herbicides penoxsulam, bispyribac-sodium, or nicosulfuron, it was also resistant to the other two. Four *ALS* mutations were characterized in association with the different patterns of cross-resistance in *E. crus-galli*. The populations surviving imazethapyr, penoxsulam, bispyribac-sodium, and nicosulfuron application had the *ALS* mutation Trp574Leu. The populations surviving only imazethapyr had one of the three following mutations: Ala122Thr, Ala205Ans, or Ser653Asn. We could link cross-resistance to each mutation, but not resistance level, because different biotypes carrying a single mutation Trp574Leu or Ser653Asn had different RF. Twenty percent of populations were still sensitive to all ALS-inhibitors sprayed, which reinforces that ALS herbicides are still important to manage *E. crus-galli* and other susceptible weed species in rice fields. The higher imazethapyr resistance frequency compared to the other three herbicides correlates with the adoption of rice cultivars resistant to imidazolinone herbicides (Clearfield® rice) that have been cultivated in Brazil since the middle 2000’s (Avila et al. 2021). The recurrent spray of imazethapyr in Clearfield® rice for about 20 years has been selecting *E. crus-galli ALS* mutations to survive imazethapyr. The use of penoxsulam and bispyribac-sodium before and concomitantly with imazethapyr in Clearfield® rice also selects mutations associated with these herbicides and imazethapyr. Penoxsulam and bispyribac-sodium use has been reduced since the availability of Clearfield® rice because they are not effective on weedy rice. Our results show that penoxsulam and bispyribac-sodium are still effective in controlling 52% of the biotypes evaluated (32% resistant only to imazethapyr + 20% susceptible) which highlights that some ALS-inhibitor active ingredients are still valuable to control *E. crus-galli*. Research papers and scientific communication sometimes generalize the resistance to one active ingredient as resistance to “ALS-inhibitors”, which is not always true and indirectly discourages the applications of the other ALS-inhibitor active ingredients. In a scenario where *Echinochloa* spp. have been evolving resistance to most of the rice selective herbicides, and few or no options are still effective in several rice fields, the understanding of *ALS* mutation cross-resistance patterns is important to design a rational ALS chemical group rotation.

There are several studies in *E. crus-galli* reporting *ALS* mutations and cross-resistance patterns, but the continuous understanding of cross-resistance patterns is important because they may change in different populations of the same weed species. For example, the mutation Ala122Thr reduced the ALS enzyme sensitivity to imazamox and penoxsulam in a previous study (Riar et al. 2013; Riar et al. 2012), while in our results the population carrying the same mutation had resistance only to imazethapyr. The complexity of cross-resistance patterns observed in *E. crus-galli* and other species supports the continuous necessity of studies diving into the target site resistance mechanism. It is important to emphasize the different resistance levels may also result from the range of doses used, environmental effect, plant development stage, moment of application, and subgenome location.

The resistance level or cross-resistance pattern of mutations is influenced by the key factors. *1) Position of the mutation*: amino acid replacements in different locations may result in resistance to different herbicides from the same mechanism of action; for example, Trp574Leu confers resistance to all ALS-inhibitors, while mutations in other positions provide resistance to one or more herbicides, as shown in our study. *2) Amino acid replacement:* different amino acids in the same position may change the affinity to different herbicides, such as Ala122Asn showing cross-resistance to four ALS-inhibitors, while our Ala122Thr only to one, and our Trp574Leu showed resistance to imidazolinones chemical group while Trp574Arg showed sensitivity to imidazolinones (Panozzo et al. 2017; Fang et al. 2019). *3) Ploidy level and number of alleles mutated*: the presence of a mutation in one homoeologous gene in a polyploid has a dilution effect in the pool of total enzymes reducing the resistance level in polyploids when compared to diploids (Yu et al. 2013). For example, the mutation Ser653Asn located only in subgenome A conferred low-resistance level in our hexaploid *E. crus-galli* populations, but showed stronger resistance in the diploids weedy rice (Ruzmi et al. 2020), *Amaranthus tuberculatus* (Patzoldt and Tranel 2007), and *A. hybridus* (Whaley et al. 2006). In addition, the presence of the Ser653Asn mutation in both BD subgenomes of wheat conferred higher imazamox resistance than the mutation in only B or D subgenomes (Hanson et al. 2006). In the same way, diploids with more than one copy of the herbicide target encoding gene in the genome show higher resistance when all genes carry a mutation; for example, soybean carrying the Pro197Ser and Trp574Leu in *ALS1* and *ALS2*, respectively, showed higher ALS-inhibitors resistance than the isolated mutations (Walter et al. 2014). Homozygosity or heterozygosity works in the same way. *4) Species/population*: different species or populations of the same species have different innate or selected increased ability to detoxify herbicides before they reach the target site, which could explain why a mutation confers different resistance patterns or resistance levels than expected. For example, the *E. crus-galli ALS* mutation Ala122Thr reduced the enzyme sensitivity to penoxsulam (Riar et al. 2013), but the population with the same mutation in our study was susceptible to penoxsulam. The findings in our study allow us to add one more factor that does not affect the cross-resistance pattern but strongly affects the resistance level, the *5) subgenome location of the mutation*: polyploid species may have the herbicide target encoding gene mutated in the subgenome that displays dominance, increasing its contribution in the pool of total ALS enzymes, making it more resistant. In addition, findings in polyploid species about the absence of *ALS* gene mutations (Carranza et al. 2023) or about the identification of one mutation without searching for sequencing in all subgenomes should be reconsidered because hidden mutations may have obscured the obtained results.

Knowledge of herbicide resistance in polyploid species has recently expanded with the generation of genomic resources, unraveling the sequence of multiple herbicide target-encoding genes. We identified that *ALS* mutations in *E. crus-galli* are prevalent in subgenome A, the least expressed, while two biotypes showed a mutation in *ALS* from subgenome C, which is the most expressed. The biotype with Trp574Leu mutation in subgenome C is more resistant to ALS-inhibitors than biotypes with the same mutation in subgenome A, with a clear impact on resistance level depending on which subgenome the mutations are located. The interaction of location × subgenome dominance advances the target site resistance understanding adding one more factor driving the herbicide resistance level in *E. crus-galli*. In addition, we identified four *ALS* mutations in *E. crus-galli* in Southern Brazil, and three of them could be effectively inhibited by ALS-inhibiting herbicides other than imidazoline chemical group. The rotation of ALS-inhibitors chemical groups could be a strategy to control the imidazoline-resistant populations carrying Ala122Thr, Ala205Asn, and Ser653Asn mutations.

## Supporting information description

**Suppl. Fig. 1**. Alignment of the whole *ALS* gene from the three subgenomes showing the SNPs that differentiate them and specific primers targeting each one.

**Suppl. Table 1.** Doses of four ALS-inhibitors utilized for the dose-response curves

**Suppl. Table 2.** Primers utilized in copy number variation and relative gene expression assays.

**Suppl. Table 3.** Screening result of 100 *Echinochloa* populations from Southern Brazil with four ALS-inhibitor chemical groups.

## Declaration of competing interest

The authors have no conflicts of interest to declare.

## Data availability

All the data are available in the manuscript and supplementary material. Additional data will be made available upon request.

## Notes

### Competing Interest Statement

The authors have declared no competing interest.

