## Supporting information for "Subgenome dominance of *ALS* target site mutations impacts herbicide resistance in allohexaploid *Echinochloa crus-galli*"

### SUPPLEMENTARY MATERIAL

```
ALS-A      ATGGCCACGACCGCCGCCGCCACCGCCGCCACGGCGGCCGCCGCGCTCACCGGCGCCACC  60
ALS-B      ATGGCCACGACCGCCGCCGCCACCGCCGCCACGGCGGCCGCCGCGCTCACCGGCGCCACC  60
ALS-C      ATGGCCACGACCGCCGCCGCCACCGCCGCCACGGCGGCCGCCGCGCTCACCGGCGCCACC  60
           ************************************************************

ALS-A      ACCGCCGCGCCCAGGCCGAGCCGCCACGGTTACTCCGCGGCGGCCGCCCGCCGCGCCGCG  120
ALS-B      ACCGCCGCGCCCAGGCCGAGCCGCCACGGTTACTCCGCGGCGGCCGCCCGCCGCGCCGCG  120
ALS-C      ACCGCCGCGCCCAGGCCGAGCCGCCACGGTTACTCCGCGGCGGCCGCCCGCCGCGCCGCG  120
           ************************************************************

ALS-A      CCCATCCGGTGCTCCGCGGCGTCGCCGGCCACGGCCACGGCTCCCCCGGCCACCCCGCTC  180
ALS-B      CCCATCCGGTGCTCCGCGGCGTCGCCGGCCACGGCCACGGCTCCCCCGGCCACCCCGCTC  180
ALS-C      CCCATCCGGTGCTCCGCGGCGTCGCCGGCCACGGCCACGGCTCCCCCGGCCACCCCGCTC  180
           ************************************************************

ALS-A      CGACCGTGGGGCCCCACCGAGCCGCGCAAGGGCGCCGACATCCTCGTCGAGGCCCTCGAG  240
ALS-B      CGACCGTGGGGCCCCACCGAGCCGCGCAAGGGCGCCGACATCCTCGTCAAGGCCCTCGAG  240
ALS-C      CGACCGTGGGGCCCCACCGAGCCGCGCAAGGGCGCCGACATCCTCGTCGAGGCCCTCGAG  240
           ************************************************ ***********

ALS-A      CGCTGCGGCGTCCGCGACATCTTCGCCTACCCCGGCGGCGCCTCCATGGAGATCCACCAG  300
ALS-B      CGCTGCGGCGTCCGCGACGTCTTCGCCTACCCCGGCGGCGCCTCCATGGAGATCCACCAG  300
ALS-C      CGCTGCGGCGTCCGCGACATCTTCGCCTACCCCGGCGGCGCCTCCATGGAGATCCACCAG  300
           ****************** *****************************************

ALS-A      GCGCTCACCCGCTCCCCCGTCATCGCCAACCACCTCTTCCGCCACGAGCAAGGGGAGGCC  360
ALS-B      GCGCTCACCCGCTCCCCCGTCATCGCCAACCACCTCTTCCGCCACGAGCAAGGGGAGGCC  360
ALS-C      GCGCTCACCCGCTCCCCCGTCATCGCCAACCACCTCTTCCGCCACGAGCAGGGGGAGGCC  360
           ************************************************** *********

ALS-A      TTCGCCGCCTCCGGGTTCGCGCGCTCGTCCGGCCGCGTCGGCGTCTGCGTCGCCACCTCG  420
ALS-B      TTCGCCGCCTCCGGGTTCGCGCGCTCGTCCGGCCGCGTCGGCGTCTGCGTCGCCACCTCG  420
ALS-C      TTCGCCGCCTCCGGCTTCGCGCGCTCGTCCGGCCGCGTCGGCGTCTGCGTCGCCACCTCG  420
           ************** *********************************************

ALS-A      GGCCCCGGCGCCACCAACCTCGTCTCCGCGCTCGCCGACGCGCTGCTCGACTCCATCCCC  480
ALS-B      GGCCCCGGCGCCACCAACCTCGTCTCCGCGCTCGCCGACGCGCTGCTCGACTCCATCCCC  480
ALS-C      GGCCCCGGCGCCACCAACCTCGTCTCCGCGCTCGCCGACGCGGTGCTCGACTCCATCCCC  480
           ****************************************** *****************

ALS-A      ATGGTCGCCATCACCGGCCAGGTGCCCCGCCGCATGATCGGCACCGACGCCCTCCAGGAG  540
ALS-B      ATGGTCGCCATCACCGGCCAGGTGCCCCGCCGCATGATCGGCACCGACGGCTTCCAGGAG  540
ALS-C      ATGGTCGCCATCACCGGCCAGGTGCCCCGCCGCATGATCGGCACCGACGCCTTCCAGGAG  540
           ************************************************* * ********

ALS-A      ACGCCAATCGTCGAGGTCACCCGCTCAATCACCAAGCACAACTACCTCGTCCTCGACATC  600
ALS-B      ACGCCAATCGTCGAGGTCACCCGCCCCATCACCAAGCACAACTACCTCGTCCTCGACATC  600
```

```
ALS-C       ACGCCAATCGTCGAGGTCACCCGCTCCATCACCAAGCACAACTACCTCGTCCTCGACATC  600
            ************************ * *********************************

ALS-A       GACGACATCCCCCGCGTCGTGCAGGAGGCGTTCTTCCTCGCCTCCTCTGGCCGACCGGGG  660
ALS-B       GACGACATCCCCCGCGTCATACAGGAGGCCTTCTTCCTCGCCTCCTCTGGCCGGCCCGGG  660
ALS-C       GACGACATCCCCCGCGTCATACAGGAGGCGTTCTTCCTCGCCTCCTCTGGCCGGCCGGGG  660
            ****************** * ******** *********************** ** ***

ALS-A       CCGGTGCTCGTCGACATCCCCAAGGACATCCAGCAGCAGATGGCCGTGCCGGTCTGGAAC  720
ALS-B       CCGGTGCTCGTCGACATCCCCAAGGACATCCAGCAGCAGATGGCCGTGCCGGTCTGGAAC  720
ALS-C       CCGGTGCTCGTCGACATCCCCAAGGACATCCAGCAGCAGATGGCCGTGCCCGTCTGGAAC  720
            ************************************************** *********

ALS-A       ACGCCCATGAGTCTGCCGGGGTACATTGCGCGCCTGCCCAAGCCTCCGGCAACTGAATTG  780
ALS-B       ACGCCCATGAGTCTGCCGGGGTACATTGCGCGCCTGCCCAAGCATCCGGCAACTGAATTG  780
ALS-C       ACGCCCATGAGTCTGCCGGGGTACATTGCGCGCCTGCCCAAGCCTCCGGCAACTGAATTG  780
            ******************************************* ****************

ALS-A       CTTGAGCAGGTGCTGCGTCTTGTTGGTGAGTCACGGCGCCCTGTTCTTTATGTTGGTGGT  840
ALS-B       CTTGAGCAGGTGCTGCGTCTTGTTGGTGAGTCGCGGCGCCCTGTTCTTTATGTTGGTGGT  840
ALS-C       CTTGAGCAGGTGCTGCGTCTTGTTGGTGAGTCACGGCGCCCTGTTCTTTATGTTGGTGGT  840
            ******************************** ***************************

ALS-A       GGCTGCGCTGCATCCGGTGAGGAGCTGTGCCGCTTTGTGGAGATGACCGGAATCCCAGTG  900
ALS-B       GGTTGCGCTGCATCCGGTGAGGAGCTGCGCCGCTTTGTGGAGATGACCGGAATCCCAGTG  900
ALS-C       GGTTGCGCTGCATCCGGTGAGGAGCTGCGCCGCTTTGTGGAGATGACCGGAATCCCAGTG  900
            ** ************************ ********************************

ALS-A       ACAACTACTCTGATGGGCCTTGGCAACTTCCCCAGTGATGACCCACTGTCTCTGCGCATG  960
ALS-B       ACAACTACTCTAATGGGCCTTGGTAACTTCCCCAGTGATGATCCACTGTCTCTGCGCATG  960
ALS-C       ACAACTACTCTGATGGGCCTTGGCAACTTCCCCAGTGATGACCCACTGTCTCTGCGCATG  960
            *********** *********** ***************** ******************

ALS-A       CTTGGTATGCACGGTACTGTATATGCAAATTATGCAGTGGATAAGGCCGACCTGTTGCTG  1020
ALS-B       CTTGGTATGCACGGTACTGTATATGCAAATTATGCAGTGGATAAGGCCGACCTGTTGCTG  1020
ALS-C       CTCGGTATGCACGGTACTGTATATGCAAATTATGCAGTGGATAAGGCCGACCTGTTGCTG  1020
            ** *********************************************************

ALS-A       GCGTTTGGTGTGCGGTTCGATGATCGCGTGACAGGAAAAATTGAGGCTTTTGCAAGCAGG  1080
ALS-B       GCATTTGGTGTGCGGCTCGATGATCGTGTGACAGGGAAAATTGAGGCTTTTGCAAGCAGG  1080
ALS-C       GCATTTGGTGTGCGGTTCGATGATCGTGTGACAGGGAAAATTGGGGCTTTTGCAGGCAGG  1080
            ** ************ ********** ******** ******* ********** *****

ALS-A       GCCAAGATTGTGCACATTGATATTGATCCAGCTGAGATTGGCAAGAACAAGCAGCCACAT  1140
ALS-B       GCCAAGATTGTGCACATTGATATTGATCCAGCTGAGATTGGCAAGAACAAGCAGCCACAT  1140
ALS-C       GCCAAGATTGTGCACATTGATATTGATCCAGCTGAGATTGGCAAGAACAAGCAGCCACAT  1140
            ************************************************************

ALS-A       GTGTCCATCTGTGCGGATGTCAAGCTTGCTTTGCAGGGCATGAATGCTCTTCTGGAAGGA  1200
ALS-B       GTGTCCATCTGTGCGGATGTCAAGCTTGCTTTGCAGGGCATGAATGCTCTTCTGGAAGGA  1200
ALS-C       GTGTCCATCTGTGCGGATGTCAAGCTTGCTTTGCAGGGCATGAATGCTCTTCTGGAAGGA  1200
```

```
           ************************************************************

ALS-A      ATCATATCAAAGAAGAGTTTTGACTTTGGCTCATGGCACGATGAGTTGGATCAGCAGAAG  1260
ALS-B      ATCATATCAAAGAAGAGTTTTGACTTTGGCTCATGGCAAGATGAGTTGGATCAGCAGAAG  1260
ALS-C      ATCATATCAAAGAAGAGTTTTGACTTTGGCTCATGGCAGGATGAGTTGGATCAGCAGAAG  1260
           ************************************** *********************

ALS-A      AGGGAATTCCCCCTGGGGTACAAAACTTTCGATGAGGAGATTCAGCCACAGTATGCTATC  1320
ALS-B      AGGGAATTCCCCCTGGGGTACAAAACTTTCGATGAGGAGATTCAGCCACAGTATGCTATC  1320
ALS-C      AGGGAATTCCCCCTGGGGTACAAAACTTTTGATGAGGAGATTCAGCCACAGTATGCTATC  1320
           ***************************** ******************************

ALS-A      CAGGTTCTGGATGAGCTGACGAAAGGGGAGGCCATCATTGCCACTGGTGTTGGGCAGCAC  1380
ALS-B      CAGGTTCTGGATGAGCTGACCAAAGGGGAGGCCATCATTGCCACTGGTGTTGGGCAGCAC  1380
ALS-C      CAGGTTCTGGATGAGCTGACGAAAGGGGAGGCCATCATTGCCACTGGTGTTGGGCAACAC  1380
           ******************** *********************************** ***

ALS-A      CAGATGTGGGCGGCACAGTACTACACTTACAAGCGGCCAAGGCAGTGGTTGTCTTCAGCT  1440
ALS-B      CAGATGTGGGCGGCACAGTACTACACTTACAAGCGGCCAAGGCAGTGGTTGTCTTCAGCT  1440
ALS-C      CAGATGTGGGCGGCACAGTACTACACTTACAAGCGACCAAGGCAGTGGTTGTCTTCAGCT  1440
           *********************************** ************************

ALS-A      GGTCTTGGGGCTATGGGATTTGGTTTACCAGCTGCTGCTGGTGCTGCTGTGGCCAACCCA  1500
ALS-B      GGTCTTGGGGCTATGGGATTTGGTTTGCCAGCTGCTGCTGGTGCTGCTGTGGCCAACCCA  1500
ALS-C      GGTCTTGGAGCTATGGGATTTGGTTTGCCGGCTGCTGCTGGTGCTGCTGTGGCGAACCCA  1500
           ******** ***************** ** *********************** ******

ALS-A      GGTGTTACAGTTGTTGACATCGATGGGGATGGCAGCTTCCTCGTGAACATTCAGGAGTTG  1560
ALS-B      GGTGTTACAGTTGTTGACATCGATGGGGATGGCAGCTTCCTCATGAACATTCAGGAGTTG  1560
ALS-C      GGTGTTACAGTTGTTGACATCGATGGGGATGGCAGCTTCCTCATGAACATTCAGGAGTTG  1560
SubB_F                                                            AGTTG
           ****************************************** *****************

ALS-A      GCTATGATCCGCATTGAGAACCTCCCAGTGAAGGTCTTTGTGCTAAACAACCAACACCTG  1620
ALS-B      GCTATGATCCGCATCGAGAACCTCCCAGTGAAGGTCTTTGTGCTAAACAACCAACACCTG  1620
ALS-C      GCTATGATCCGCATTGAGAACCTCCCAGTGAAGGTCTTTGTGCTAAACAACCAACACCTT  1620
SubB_F     GCTATGATCCGCATC
           ************** ********************************************

ALS-A      GGGATGGTGGTGCAGTGGGAGGACAGATTCTACAAGGCCAACCGAGCACACACATACTTG  1680
ALS-B      GGTATGGTGGTGCAGTGGGAGGACAGATTCTACAAGGCCAACAGAGCACACACATACTTG  1680
ALS-C      GGGATGGTGGTGCAGTGGGAGGACAGATTCTACAAGGCCAACCGAGCACATACATACTTG  1680
SubA_F                                                GAGCACACACATACTTG
           ** *************************************** ******* *********

ALS-A      GGGAATCCAGAGAATGAGAGCGAGATATATCCGGATTTCGTGACGATTGCCAAAGGATTC  1740
ALS-B      GGGAACCCAGAGAATGAGAGCGAGATATATCCGGATTTTGTAACGATTGCCAAAGGGTTC  1740
ALS-C      GGGAACCCAGATAATGAGAGCGAGATATATCCGGATTTCGTGACCATTGCCAAAGGTTTT  1740
SubA_F     GGGCAT
SubC_F                            GATATATCCGGATTTCGTGACC
           ***** ***** ************************** ** ** *********** **
```

```
ALS-A      AACATTCCAGCAGTCCGTGTGACAAAGAAGAGCGAAGTCCGTGCAGCAATTAAGAAGATG  1800
ALS-B      AACATTCCAGCAGTCCGTGTGACAAAGAAGAGCGAAGTACGTGCAGCAATCAAGAAGATG  1800
ALS-C      AACATTCCAGCGGTCCGTGTGACAAAGAAGAGCGAAGTACGTGCAGCAATCAAGAAGATG  1800
           *********** ************************** *********** *********

ALS-A      CTCGAGACTCCAGGGCCGTACCTGTTGGATATCATTGTCCCGCACCAGGAACATGTGTTG  1860
ALS-B      CTCGAGACTCCAGGGCCGTACCTGTTGGATATCATTGTCCCGCACCAGGAACATGTGTTG  1860
ALS-C      CTCGAGACTCCAGGGCCATACCTGTTGGATATCATTGTCCCGCACCAGGAACATGTGTTG  1860
           ***************** ******************************************

ALS-A      CCTATGATCCCGAGCGGTGGCGCTTTCAAGGACATGATCCTGGATGGTGATGGCAGGACC  1920
ALS-B      CCTATGATCCCGAGCGGTGGCGCTTTCAAGGACATGATCCTGGATGGTGATGGCAGGACC  1920
ALS-C      CCTATGATCCCGAGCGGTGGCGCTTTCAAGGACATGATCCTGGATGGTGATGGCAGGACC  1920
           ************************************************************

ALS-A      GTGTATTGA  1929
ALS-B      GTGTATTGA  1929
ALS-C      GTGTATTGA  1929
           *********
```

Suppl. Fig. 1. Alignment of the whole *ALS* gene from three subgenomes showing the SNPs that differentiate them and specific primers targeting each one. The sequences are from Genbank: MH013494.1 (*ALS*-SubA), MG188321.1 (*ALS*-SubB), and MH013493.1 (*ALS*-SubC). Asterisk (*) indicates the same nucleotide in all three subgenomes. Absence of * indicates there is at least one different nucleotide between them.

Suppl. Table 1. Doses of four ALS-inhibitors utilized for the dose-response curve with *E. crus-galli* biotypes with different *ALS* mutations

| Biotypes | Doses | Label rate range |
|---|---|---|
| | Imazethapyr | |
| MOSTS-01 and CAPL-01 | 0; 3.312; 6.625; 13.25; 26.5; 53; 106; 212 g $ha^{-1}$ | 0 - 2x |
| ARRGR-01 and PALMS-01 | 0; 13.25; 26.5; 53; 106; 212; 424; 848 g $ha^{-1}$ | 0 - 8x |
| BAGÉ-01, SANTPAT-01, 423, CAMAQ-01, SAOJER-01, and CAPV-03 | 0; 53; 106; 212; 424; 848; 1,696; 3.392 g $ha^{-1}$ | 0 - 32x |
| | Penoxsulam | |
| MOSTS-01 and CAPL-01 | 0; 0.6; 3.75; 7.5; 15; 30; 60; 120 g $ha^{-1}$ | 0 - 2x |
| ARRGR-01, PALMS-01, BAGÉ-01, and SANTPAT-01 | 0; 3.75; 7.5; 15; 30; 45; 60; 120 g $ha^{-1}$ | 0 - 2x |
| 423 | 0; 3.75; 7.5; 15; 30; 60; 120; 240 g $ha^{-1}$ | 0 - 4x |
| CAPV-03 | 0; 15; 30; 60; 120; 240; 480; 960 g $ha^{-1}$ | 0 - 16x |
| CAMAQ-01 and SAOJER-01 | 0; 20; 60; 180; 540; 1,620; 4,860; 14,580 g $ha^{-1}$ | 0 - 243x |
| | Bispyribac-sodium | |
| MOSTS-01 and CAPL-01 | 0; 0.5; 3.125; 6.25; 12.5; 25; 50; 100 g $ha^{-1}$ | 0 - 2x |
| ARRGR-01, PALMS-01, BAGÉ-01, and SANTPAT-01 | 0; 6.25; 12.5; 25; 37.5; 50; 100; 200 g $ha^{-1}$ | 0 - 4x |
| 423 | 0; 3.125; 6.25; 12.5; 25; 50; 100; 200 g $ha^{-1}$ | 0 - 4x |
| CAMAQ-01, SAOJER-01, and CAPV-03 | 0; 25; 50; 100; 200; 400; 800; 1,600 g $ha^{-1}$ | 0 - 32x |
| | Nicosulfuron | |
| MOSTS-01 and CAPL-01 | 0; 0.6; 3.75; 7.5; 15; 30; 60; 120 g $ha^{-1}$ | 0 - 2x |
| ARRGR-01, PALMS-01, BAGÉ-01, and SANTPAT-01 | 0; 7.5; 15; 30; 45; 60; 120; 240 g $ha^{-1}$ | 0 - 4x |
| 423 | 0; 3.75; 7.5; 15; 30; 60; 120; 240 g $ha^{-1}$ | 0 - 4x |
| CAMAQ-01, SAOJER-01, and CAPV-03 | 0; 30; 60; 120; 240; 480; 960; 1,920 g $ha^{-1}$ | 0 - 32x |

Suppl. Table 2. Primers utilized as reference in the copy number variation and relative gene expression assays.

| | Primer | Sequence (5' - 3') | Fragment size | Annealing T°C | Reference |
|---|---|---|---|---|---|
| GAPDH | GAPDH_F | TGGAATTGCTTTGAACGACA | 99 bp | 60 | (Iwakami et al., 2014) |
| | GAPDH_R | AAGATGTGGCGGATCAGGT | | | |
| eIF4B | eIF4B_F | GGGAAGTGATTTTGCAGGAG | 190 bp | 60 | (Iwakami et al., 2014) |
| | eIF4B_R | GAGGCTTGGTCAGAACCATC | | | |
| RUB | RUB_F | GGAGTATGAAACCAAGGATACTG | 100 bp | 60 | (Duhoux & Délye, 2013) |
| | RUB_R | GTTGTCCATGTACCAGTAGAAGA | | | |
| 18S | 18S_F | GTGACGGAGAATTAGGGTTC | 100 bp | 60 | (Duhoux & Délye, 2013) |
| | 18S_R | TGTCAGGATTGGGTAATTTG | | | |
| 28S | 28S_F | CTGATCTTCTGTGAAGGGT | 200 bp | 60 | |
| | 28S_R | TGATAGAACTCGTAATGGGC | | | |

Suppl. Table 3. Screening of 100 *Echinochloa* populations from Southern Brazil with four ALS-inhibitor chemical groups: imazethapyr, penoxsualm, bispyribac-sodium, and nicosulfuron from imidazolinones, triazolopyrimidine – type 2, pyrimidinylbenzoates, and sulfonylureas, respectively. Efficacy of control 28 days after treatment (0% = no injuries; 100% = dead)

| ID | Municipality | Imazethapyr | Penoxsulam | Bispyribac-sodium | Nicosulfuron |
|---|---|---|---|---|---|
| 1 | CAMAQUÃ | 25.00 | 50.00 | 30.00 | 10.00 |
| 2 | TAPES | 21.67 | 100.00 | 100.00 | 95.00 |
| 3 | TAPES | 30.00 | 100.00 | 100.00 | 85.00 |
| 4 | TAPES | 75.00 | 81.67 | 58.33 | 75.00 |
| 5 | TAPES | 100.00 | 100.00 | 100.00 | 93.33 |
| 6 | TAPES | 48.33 | 23.33 | 40.00 | 50.00 |
| 7 | VENANCIO AIRES | 26.67 | 3.33 | 41.67 | 11.67 |
| 8 | SANTA CRUZ | 23.33 | 43.33 | 56.67 | 41.67 |
| 9 | SANTA CRUZ | 60.00 | 30.00 | 20.00 | 48.33 |
| 10 | SÃO PEDRO | 20.00 | 0.00 | 55.00 | 18.33 |
| 11 | SÃO PEDRO | 46.67 | 100.00 | 100.00 | 93.33 |
| 12 | SÃO PEDRO | 50.00 | 51.67 | 26.67 | 30.00 |
| 13 | AGUDO | 40.00 | 100.00 | 100.00 | 100.00 |
| 14 | PARAISO DO SUL | 41.67 | 0.00 | 31.67 | 10.00 |
| 15 | MOSTARDAS | 33.33 | 0.00 | 21.67 | 10.00 |
| 16 | CAPÃO DO LEAO | 50.00 | 100.00 | 100.00 | 100.00 |
| 17 | CAPÃO DO LEAO | 93.33 | 100.00 | 100.00 | 100.00 |
| 18 | CAPÃO DO LEAO | 18.33 | 100.00 | 95.00 | 93.33 |
| 19 | PIRATINI | 65.00 | 65.00 | 36.67 | 25.00 |
| 20 | PIRATINI | 43.33 | 0.00 | 25.00 | 20.00 |
| 21 | HULHA NEGRA | 53.33 | 21.67 | 53.33 | 50.00 |
| 22 | ACEGUÁ | 50.00 | 50.00 | 31.67 | 20.00 |
| 23 | BAGE | 50.00 | 100.00 | 91.67 | 100.00 |
| 24 | DOM PEDRITO | 48.33 | 15.00 | 25.00 | 15.00 |
| 25 | DOM PEDRITO | 31.67 | 21.67 | 36.67 | 10.00 |
| 26 | DOM PEDRITO | 21.67 | 100.00 | 100.00 | 90.00 |
| 27 | DOM PEDRITO | 56.67 | 0.00 | 15.00 | 0.00 |
| 28 | CHARQUEADAS | 60.00 | 100.00 | 100.00 | 100.00 |
| 29 | PANTANO GRANDE | 75.00 | 100.00 | 100.00 | 95.00 |
| 30 | RIO PARDO | 56.67 | 100.00 | 85.00 | 90.00 |
| 31 | MASSAMBARÁ | 100.00 | 100.00 | 100.00 | 100.00 |
| 32 | CAMAQUÃ | 50.00 | 46.67 | 30.00 | 35.00 |
| 33 | URUGUAIANA | 75.00 | 100.00 | 100.00 | 100.00 |
| 34 | URUGUAIANA | 100.00 | 100.00 | 100.00 | 100.00 |
| 35 | URUGUAIANA | 53.33 | 33.33 | 31.67 | 20.00 |
| 36 | ARROIO GRANDE | 100.00 | 100.00 | 100.00 | 100.00 |
| 37 | CAMAQUÃ | 48.33 | 100.00 | 100.00 | 100.00 |
| 38 | SANTA VITORIA DO PALMAR | 20.00 | 23.33 | 45.00 | 10.00 |

| | | | | | |
|---|---|---|---|---|---|
| 39 | URUGUAIANA | 46.67 | 35.00 | 33.33 | 16.67 |
| 40 | SANTA VITORIA DO PALMAR | 100.00 | 100.00 | 100.00 | 100.00 |
| 41 | TUBARÃO | 26.67 | 20.00 | 3.33 | 15.00 |
| 42 | IÇARA | 20.00 | 20.00 | 43.33 | 10.00 |
| 43 | MOSTARDAS | 35.00 | 26.67 | 10.00 | 1.67 |
| 44 | PELOTAS | 50.00 | 40.00 | 26.67 | 25.00 |
| 45 | ITAJAÍ | 55.00 | 53.33 | 33.33 | 70.00 |
| 46 | ARARANGUÁ | 18.33 | 20.00 | 45.00 | 0.00 |
| 47 | PELOTAS | 100.00 | 100.00 | 100.00 | 100.00 |
| 48 | CAMAQUÃ | 25.00 | 100.00 | 100.00 | 100.00 |
| 49 | CAMAQUÃ | 45.00 | 23.33 | 33.33 | 28.33 |
| 50 | CACHOEIRA DO SUL | 30.00 | 26.67 | 38.33 | 33.33 |
| 51 | RIO PARDO | 100.00 | 100.00 | 100.00 | 100.00 |
| 52 | TAPES | 32.50 | 93.33 | 100.00 | 100.00 |
| 53 | TAPES | 45.00 | 100.00 | 100.00 | 100.00 |
| 54 | VIAMAO | 100.00 | 100.00 | 100.00 | 100.00 |
| 55 | AGUDO | 43.33 | 20.00 | 45.00 | 43.33 |
| 56 | ITAQUI | 10.00 | 20.00 | 35.00 | 13.33 |
| 57 | RIO GRANDE | 23.33 | 22.50 | 66.67 | 31.67 |
| 58 | MOSTARDAS | 100.00 | 100.00 | 100.00 | 100.00 |
| 59 | CACHOEIRA DO SUL | 100.00 | 100.00 | 100.00 | 100.00 |
| 60 | MOSTARDAS | 100.00 | 100.00 | 100.00 | 100.00 |
| 61 | CACHOEIRINHA | 100.00 | 100.00 | 100.00 | 100.00 |
| 62 | CACHOEIRINHA | 100.00 | 100.00 | 100.00 | 100.00 |
| 63 | CACHOEIRINHA | 100.00 | 100.00 | 100.00 | 100.00 |
| 64 | SANTA MARIA-01 | 100.00 | 100.00 | 100.00 | 100.00 |
| 65 | ANTA GORDA-01 | 100.00 | 100.00 | 100.00 | 100.00 |
| 66 | CAPAO DO LEÃO | 100.00 | 100.00 | 100.00 | 100.00 |
| 67 | PORTO ALEGRE | 100.00 | 100.00 | 100.00 | 100.00 |
| 68 | VALE VERDE | 100.00 | 100.00 | 100.00 | 100.00 |
| 69 | BAGE | 30.00 | 100.00 | 100.00 | 100.00 |
| 70 | PALMARES DO SUL | 30.00 | 100.00 | 100.00 | 100.00 |
| 71 | MOSTARDAS | 43.33 | 50.00 | 50.00 | 20.00 |
| 72 | CAMAQUÃ | 50.00 | 70.00 | 51.67 | 36.67 |
| 73 | RIO GRANDE | 40.00 | 50.00 | 35.00 | 33.33 |
| 74 | RIO PARDO | 40.00 | 40.00 | 40.00 | 23.33 |
| 75 | PALMARES DO SUL | 63.33 | 100.00 | 100.00 | 100.00 |
| 76 | CAPOVARI DO SUL | 40.00 | 38.33 | 30.00 | 23.33 |
| 77 | SÃO JERONIMO | 10.00 | 30.00 | 0.00 | 23.33 |
| 78 | SANTO ANTONIO DA PATRULHA | 20.00 | 100.00 | 100.00 | 100.00 |
| 79 | ARROIO GRANDE | 33.33 | 100.00 | 100.00 | 100.00 |
| 80 | CACHOEIRA DO SUL | 75.00 | 100.00 | 100.00 | 100.00 |
| 81 | ARROIO GRANDE | 65.00 | 100.00 | 100.00 | 100.00 |

| | | | | | |
|---|---|---|---|---|---|
| 82 | ARROIO GRANDE | 73.33 | 100.00 | 100.00 | 100.00 |
| 83 | CACHOEIRA DO SUL | 33.33 | 100.00 | 100.00 | 100.00 |
| 84 | CANDELÁRIA | 30.00 | 100.00 | 100.00 | 100.00 |
| 85 | CANDELÁRIA | 33.33 | 100.00 | 100.00 | 100.00 |
| 86 | CACHOEIRA DO SUL | 50.00 | 52.50 | 50.00 | 56.67 |
| 87 | AGUDO | 20.00 | 100.00 | 100.00 | 100.00 |
| 88 | ARROIO GRANDE | 100.00 | 100.00 | 100.00 | 100.00 |
| 89 | CERRITO | 18.33 | 100.00 | 100.00 | 100.00 |
| 90 | CERRITO | 30.00 | 100.00 | 100.00 | 100.00 |
| 91 | MOSTARDAS | 33.33 | 96.67 | 100.00 | 100.00 |
| 92 | PIRATINI | 31.67 | 3.33 | 0.00 | 15.00 |
| 93 | PIRATINI | 25.00 | 26.67 | 18.33 | 21.00 |
| 94 | HULHA NEGRA | 20.00 | 36.67 | 25.00 | 25.00 |
| 95 | DOM PEDRITO | 6.67 | 33.33 | 45.00 | 35.00 |
| 96 | ITAQUI | 0.00 | 6.67 | 10.00 | 12.00 |
| 97 | ITAQUI | 3.33 | 5.00 | 26.67 | 18.00 |
| 98 | ITAQUI | 13.33 | 0.00 | 25.00 | 30.00 |
| 99 | URUGUAIANA | 0.00 | 0.00 | 23.33 | 14.00 |
| 100 | CACEQUI | 51.67 | 66.67 | 36.67 | 35.00 |
